# Neuronal segmentation in cephalopod arms

**DOI:** 10.1101/2024.05.29.596333

**Authors:** Cassady S. Olson, Natalie Grace Schulz, Clifton W. Ragsdale

## Abstract

The prehensile arms of the cephalopod are among these animals most remarkable features, but the neural circuitry governing arm and sucker movements remains largely unknown. We studied the neuronal organization of the adult axial nerve cord (ANC) of *Octopus bimaculoides* with molecular and cellular methods. The ANCs, which lie in the center of every arm, are the largest neuronal structures in the octopus, containing four times as many neurons as found in the central brain. In transverse cross section, the cell body layer (CBL) of the ANC wraps around its neuropil (NP) with little apparent segregation of sensory and motor neurons or nerve exits. Strikingly, when studied in longitudinal sections, the ANC is segmented. ANC neuronal cell bodies form columns separated by septa, with 15 segments overlying each pair of suckers. The segments underlie a modular organization to the ANC neuropil: neuronal cell bodies within each segment send the bulk of their processes directly into the adjoining neuropil, with some reaching the contralateral side. In addition, some nerve processes branch upon entering the NP, forming short-range projections to neighboring segments and mid-range projections to the ANC segments of adjoining suckers. The septa between the segments are employed as ANC nerve exits and as channels for ANC vasculature. Cellular analysis establishes that adjoining septa issue nerves with distinct fiber trajectories, which across two segments (or three septa) fully innervate the arm musculature. Sucker nerves also use the septa, setting up a nerve fiber “suckerotopy” in the sucker-side of the ANC. Comparative anatomy suggests a strong link between segmentation and flexible sucker-laden arms. In the squid *Doryteuthis pealeii*, the arms and the sucker- rich club of the tentacles have segments, but the sucker-poor stalk of the tentacles does not. The neural modules described here provide a new template for understanding the motor control of octopus soft tissues. In addition, this finding represents the first demonstration of nervous system segmentation in a mollusc.

## Main body

The octopus has a motor control challenge of enormous complexity^1,2^. Each of its eight arms is a muscular hydrostat, a soft-bodied structure that lacks a rigid skeleton and moves with near infinite degrees of freedom^3,4^. Moreover, the arms are packed with hundreds of chemotactile suckers which can change shape independently^5^. Even with this complexity, octopuses control behaviors effectively along the length of a single arm, across all eight arms and between suckers^6–8^. The neural circuits underlying these behaviors are unexplored with modern molecular and cellular methods.

Embedded in the octopus arm is a massive nervous system, with more neurons found distributed across the eight arms than in the brain^9,10^. Most prominent is an axial nerve cord (ANC) running down the center of every arm (Fig 1a)^2,11^. Peripheral to the ANC, there are four small intramuscular nerve cords (IMNCs), and a sucker ganglion (SG) for every sucker (Fig 1a). In the ANC, and following the characteristic invertebrate pattern, neuronal cell bodies are localized to a cell body layer (CBL) wrapping around neuropil (NP). In transverse sections, the CBL forms a horseshoe on the sucker, or oral, side of the ANC (Fig 1b). On the aboral side, or away from the suckers, there is a massive cerebrobrachial tract (CBT) connecting the arms and the brain (Fig 1b). The CBL can be divided into an aboral, or brachial, territory that is dedicated to the sensorimotor control of the arm, and an oral, or sucker, territory for the sensorimotor control of the suckers^12^ (SFig 1f, g).

**Figure 1:**
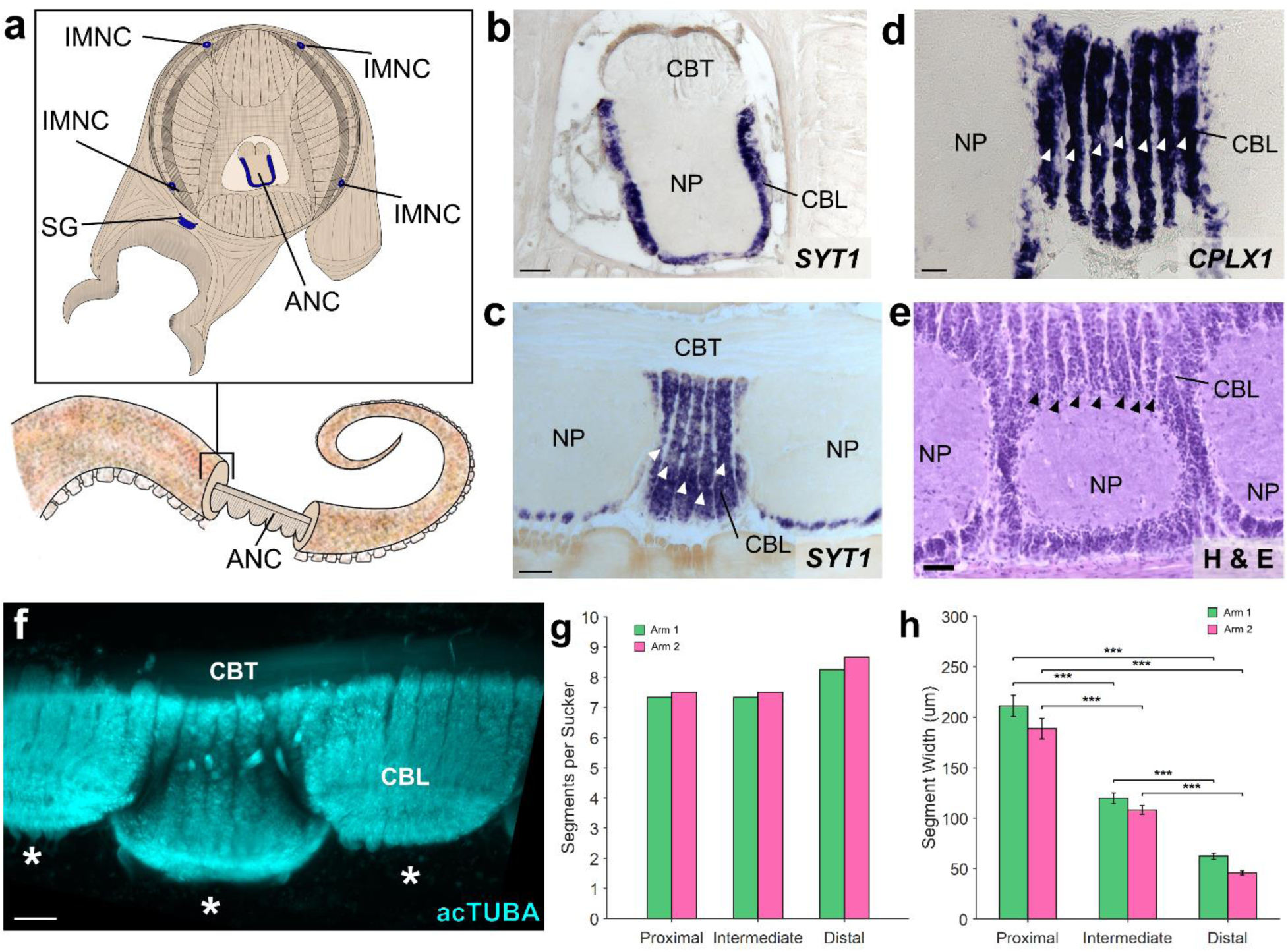
The axial nerve cord (ANC) is segmented. **a,** Transverse diagram of *O. bimaculoides* arm with the location of neuronal cell bodies outlined in blue. **b,** In situ hybridization (ISH) for *SYT1* in a transverse section of the ANC. Scale bar: 100 μm. **c-e:** Longitudinal sections through the ANC demonstrating segmentation in the cell body layer with **(c)** ISH for *SYT1*. Scale bar: 100 μm, **(d)** ISH for *CPLX1* and **(e)** hematoxylin and eosin (H&E). Scale bars: 50 μm. **f,** Whole mount immunostaining with acetylated alpha tubulin (acTUBA) of a dissected ANC. Segmentation pattern is continuous as the ANC oscillates from sucker to sucker. * denotes individual suckers. Scale bar: 100 μm. **g-h.** Quantification of segmentation down the proximal-distal axis. Arm 1 (green) was stained with acTUBA and arm 2 (pink), with H&E. **(g)** The number of segments per sucker is maintained along the proximal-distal axis (average Arm 1 = 7.64; average Arm 2 = 7.88). **(h)** Segment width decreases down the proximal distal axis. n = 48 per condition, error bars +/- sem, ***p < 0.001. CBL, cell body layer; CBT, cerebrobrachial tract; IMNC, intramuscular nerve cord; NP, neuropil; SG, sucker ganglion.

Beyond this classical segregation, further neural divisions within the ANC and in the ANC nerve trajectories for the brachial musculature and the suckers are unknown.

### Segmentation in the ANC

We found an unexpected neuronal organization in the ANC along the longitudinal axis of the arm: neuronal cell bodies are constrained to segmental columns separated by septa (Fig 1c, d, e). These segments encompass the full aboral to oral extent of the cell body layer, involving both the brachial and sucker territories (SFig 1f, g). The ANC as a structure snakes down the arm, orienting in turn to each sucker, with the segmentation pattern persisting as ANC moves from sucker to sucker (Fig 1f, SFig 1e, f, g). The arm tapers and suckers become smaller along the proximal-distal axis of the arm, as do the widths of the segments (Fig 1h, SFig 1c, d). Conversely, the cellular packing density of the segments increases, and the number of segments per sucker remains constant (average: ∼7.5; Fig 1g, see also SFig 7b, e).

We next investigated the septa between segments. The space between the CBL segments is enriched with connective tissue, labeled by Picrosirius Red staining (Fig 2a). Connective tissue pads the length of the septa, and sometimes lines the interface between the CBL and NP (SFig 2d). Vasculature also lies in the septa. Most prominent is F-actin positive vasculature that travels along the inner border of the CBL and the NP, strictly following the segmental boundaries (Fig 2B, SFig 2c). To confirm that this F-actin labeling identifies blood vessels, we flooded the vascular system with dextran-TRITC. We found an overlap with vasculature labeled by dextran and F-actin (Fig 2C, SFig 2b). Thus, as with other segmental systems such as in insect developmental compartments or vertebrate somites, multiple tissue types form in a segmental pattern along the ANC^13,14^.

**Figure 2:**
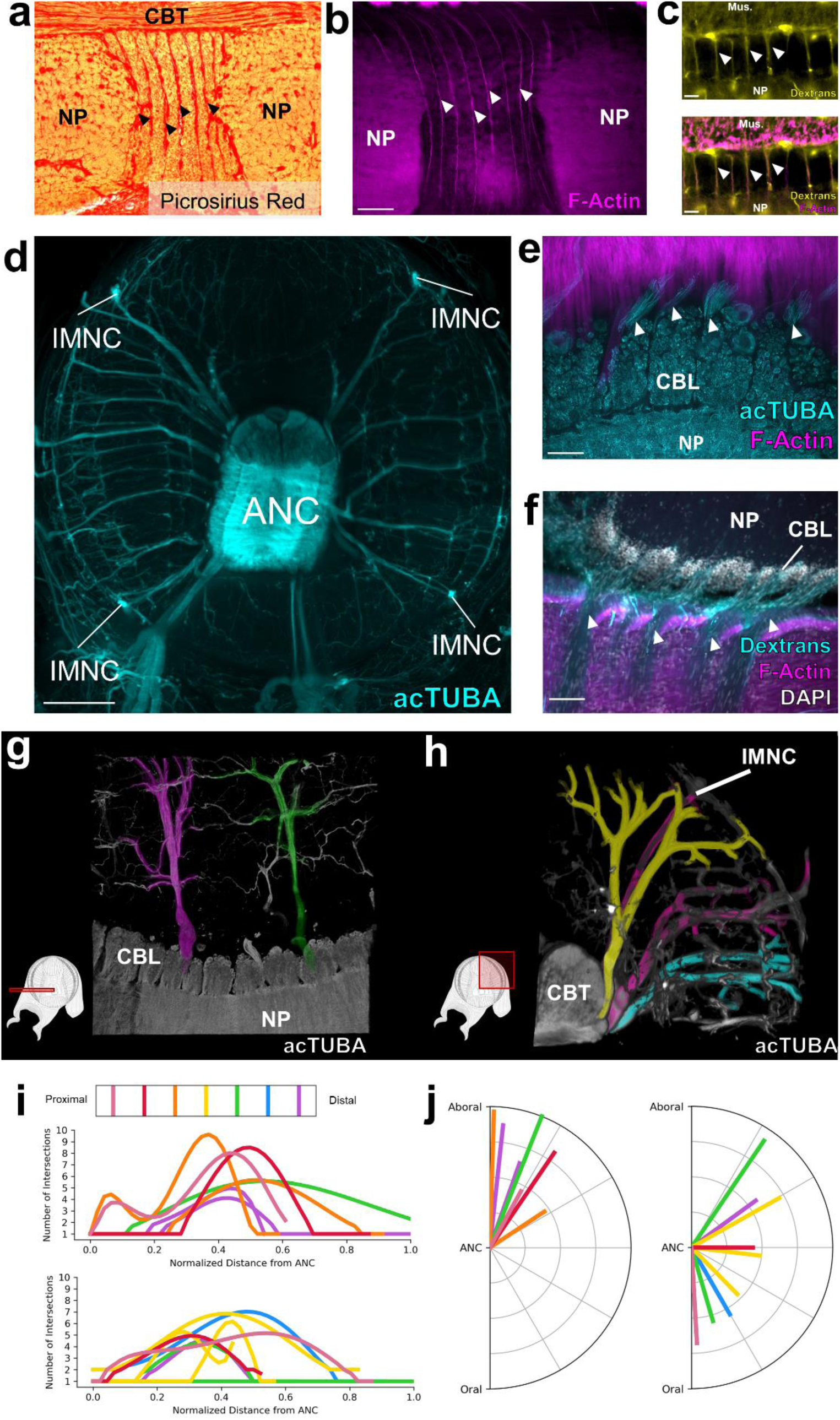
**Septa between segments are enriched for collagen, blood vessels and nerve fibers. a**, Picrosirius Red stain of an ANC longitudinal section. Red denotes the presence of collagen. **b,** F-actin (magenta) labeled by phalloidin is localized to the septa. Scale bar: 100 μm. **c,** *Top*- Trans-vascular dextran labeling demonstrates vasculature (yellow) in the septa. *Bottom*- Dextran labeling colocalizes with F-actin staining (magenta). Scale bar: 20 μm. **d,** Maximum projection of acetylated alpha-tubulin (acTUBA) whole mount immunostaining of a transverse slice. Nerve fibers exit the ANC at discrete locations. Scale bar: 500 μm. **e, f,** Longitudinal ANC sections of the brachial nerve exits in the septa. **(e)** acTUBA nerve labeling (cyan), **(f)** experimental dextran tracing of oral nerves. Injection site in central NP. Scale bars: 100 μm. **g-j** Brachial nerves exiting from neighboring septa have different targets. **(g)** Maximum projection of a horizontal whole mount stained with acTUBA. Two nerves (false colored in magenta and green) innervating similar territories of muscle are separated by multiple segments. **(h)** Maximum projection of a transverse whole mount stained with acTUBA. Three nerves (false colored with yellow, magenta, and cyan) exiting from neighboring septa target different brachial territories. **(i)** Branching profiles of nerves across multiple septa was characterized by Sholl analysis. Profiles differ from septum to septum. *Top*- Aboral nerves. *Bottom*- Central nerves. **(j)** Nerve fiber average trajectories vary across septa. *Left*- Aboral nerves. *Right*- Central nerves. Mus., brachial musculature.

### Relationship to sensorimotor circuits

Nerve fibers depart at discrete points along the oral to aboral extent of the ANC, utilizing the septa as exit points (Fig 2d, e, f). Physiological and classic neuroanatomical studies have shown that these nerves carry both sensory and motor information^15–17^. Given this result, we used molecular markers to ask whether there might be a segregation of sensory and motor territories within the CBL itself. Markers of motor neurons (*NKX6, MNX, LHX3*) are largely restricted to the lateral sides of the CBL (SFig 3a, b, c, d). Markers of primary sensory neurons (*DRGX, PIEZO*) are also found distributed throughout this territory (SFig 3e, f, g). These results indicate that unlike vertebrate spinal cord^18^, there is not a clear spatial separation of sensory and motor neurons within the ANC.

Nerves exiting the ANC nerves can be divided into the oral nerves, which innervate the sucker, and aboral and central nerves, which together innervate the brachial musculature (Fig 2e, f, SFig 4a). We asked how the ANC nerve fibers relate to the segmentation within the ANC by examining the innervation patterns of the aboral and central nerves targeting the brachial muscles. One possibility is that the same pattern of nerves is formed out between every segment. If this were the case, each CBL segment would reflect the same sensorimotor unit, simply spatially shifted down the longitudinal axis of the arm (See SFig 5b). We studied brachial nerve fiber trajectories with whole mount immunohistochemistry for acetylated alpha-tubulin (acTUBA) and in dextran tracer experiments. The average exit trajectory and branching profile were calculated with Sholl analysis to compare nerves exiting between adjoining and more distant septa (SFig 4). For both the central and aboral nerves, nerves exiting between adjoining septa exhibit different branching profiles and exit with different average trajectories (Fig 2h, i, j). Nerves with similar branching profiles and exit trajectories are separated by multiple septa (Fig 2g, SFig 4). We found, however, that nerve fibers across multiple septa collectively provide full coverage of the brachial musculature (SFig 5). Instead of each segment reflecting identical sensorimotor units, this innervation pattern predicts that adjoining segments collaborate in organizing the brachial muscle activation patterns. A somewhat similar alternating pattern has been documented in other segmented systems, such as that for the rhombomeres in the developing vertebrate hindbrain^19^.

We next interrogated the structure of the NP in the brachial territory of the ANC. Neuronal cell bodies send their projections directly into the NP adjacent to their segments, effectively partitioning the NP (Fig 3a, SFig 6b). Some of these projections extend across the midline, predicting crosstalk from segments on one side of the ANC to the other (Fig 3a, SFig 6c). In addition, as aboral and central nerve fibers enter the ANC, they branch into segments both proximal and distal to the septum of entry, leading to an overlap of the projections of nerve fibers from neighboring septa (Fig 3b). Collectively, this circuit structure provides the neural substrate for integration of multiple segments in the control of the brachial musculature (Fig 3c).

**Figure 3:**
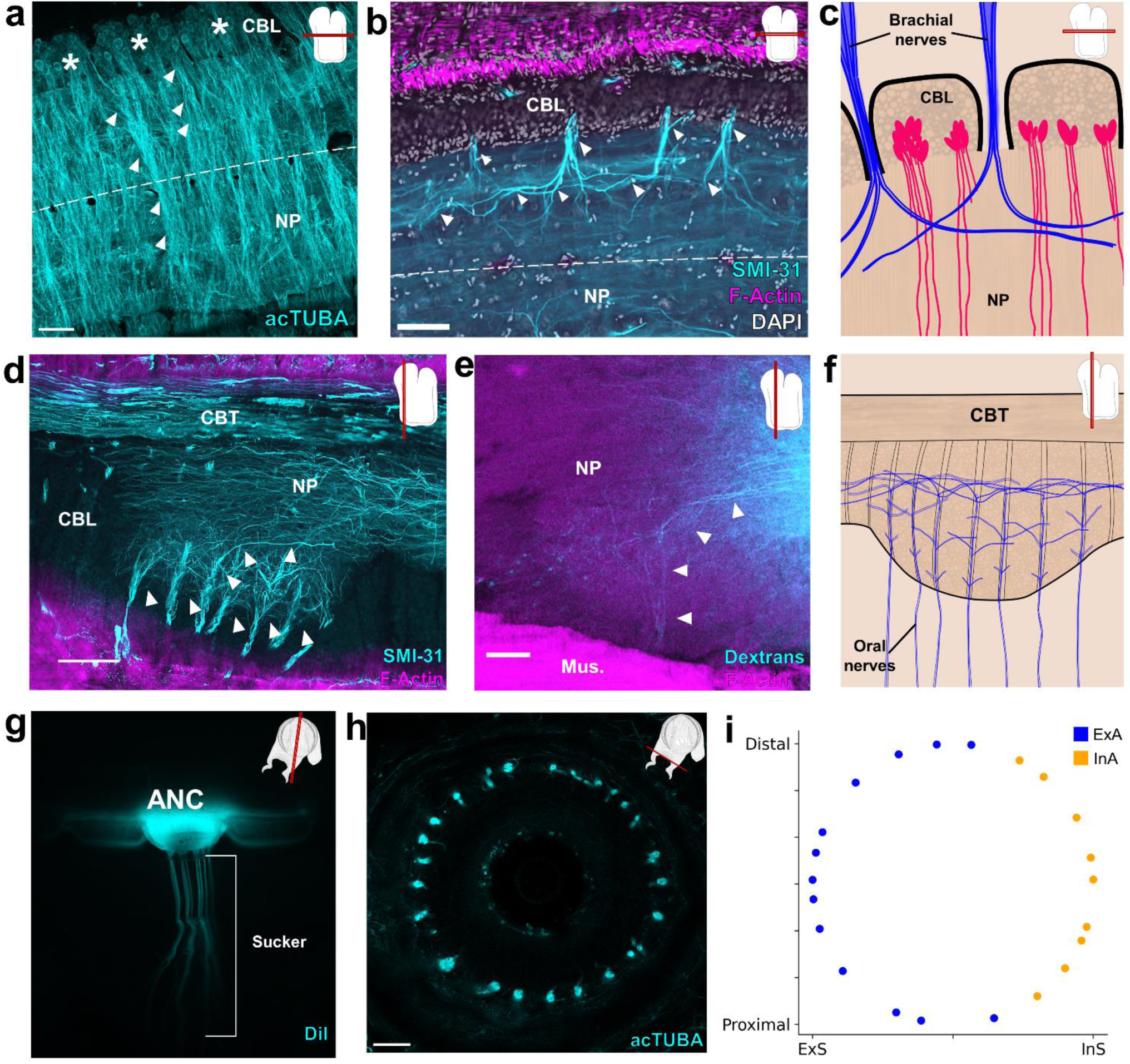
Segmental organization of the ANC neuropil. a-c, Aboral, or brachial, NP. **(a)** Maximum projection of a horizontal acTUBA wholemount (cyan) showing that the bulk of each cell body segment extends its processes into the NP of the segment. Some processes cross the midline (dashed line). * denotes individual segments. **(b)** Horizontal section immunostained with SMI-31 (cyan), phalloidin (magenta) and DAPI (gray). SMI-31-rich subpopulation of nerve fibers (cyan) branch proximally and distally upon entering the ANC. **(c)** Diagram of a horizontal section through the brachial NP. Nerve fibers (blue) pool across neurons (magenta) arranged in segments. **d-f**, Oral, or sucker, NP. **(d)** Longitudinal ANC section stained with SMI-31 (cyan) and phalloidin (magenta). NP fibers (cyan) collect into internal fascicles before exiting as oral nerves. **(e)** Longitudinal section of NP dextran tracer-deposit (cyan) demonstrating NP fibers forming a fascicle. **(f)** Diagram of NP fibers in the longitudinal plane. From oral to aboral, oral nerves enter the ANC, first showing local branching, then more extensive branching. **g-i**, Oral ANC nerves distributed around a sucker. **(g)** Longitudinal whole mount. DiI crystal (cyan) placed in a single ANC sucker enlargement targets a single sucker. **(h)** Horizontal acTUBA-stained section through a sucker demonstrates radially symmetric decoration. **(i)** Distribution of oral nerve fiber tips around a sucker. Nerves from the external ANC side (ExA) are tagged in blue; nerves from the internal side (InA), in orange. Data from a transversely imaged whole mount stained with acTUBA. The ExA covers 68% of the sucker; the InA, 32%. Scale bars: 100 μm.

The sucker is the second major target of the ANC, and we examined the relationship of oral roots innervating the sucker to the ANC segmentation. The expansion of the oral ANC opposite to each sucker was targeted for axon tracing, and, as the example illustrated in Fig 3g demonstrates, the oral nerves arising from each enlargement target a single sucker. This result was confirmed with acTUBA immunostaining (SFig 6e). After exiting the ANC in the septa, the oral roots radially tile the sucker, reaching both the muscles of the sucker and the chemotactile sensory epithelium on the rim of the sucker (Fig 3h, SFig 6f, g, h). Nerve fibers from neighboring septa target adjoining territories along the epithelium (SFig 6g). This arrangement establishes a spatial topography for the sucker in the ANC (“suckerotopy”) based on nerve exits.

We next followed the oral nerves into the ANC neuropil, where they maintain their spatial segregation, establishing an internal ANC suckerotopy (Fig 3d). As demonstrated by DiI crystal placement within a sucker, the oral nerves enter on both sides of the ANC (SFig 7f). Nerve fibers first branch locally, demonstrating short-range projections to ipsilateral nerves entering in adjoining septa (Fig 3d, SFig 6d). As the oral roots progress further into the NP, they show projections to nerves on the contralateral side (SFig 6d). Lastly, the oral roots branch to engage in mid-range projections to the ANC segments of adjoining suckers (Fig 3d, e, f). This circuitry demonstrates pathways for both intra-sucker and inter-sucker communication.

Because the oral nerves for a single sucker enter on both sides of the ANC, nerves arising from one side of the ANC connect to the internal side of the sucker (InS) and nerves arising from the other side connect to the external side of the sucker (ExS) (SFig 7a). We discovered that segments are expanded on the ExS of the ANC (ExA) as compared with the InA (SFig 7c, d). Since segments correspond to spatial territories along the sucker, this arrangement suggests that more neural territory is dedicated to the ExS. Accordingly, we found the oral roots corresponding to the ExS cover approximately 65% of the sucker, whereas the roots issued to the InS cover the remaining 35% (Fig 3i, SFig 7g, h). The sucker sensory epithelium is evenly innervated, with no obvious bias to the ExS side over the InS side (Fig 3h, SFig 6h). Consequently, the asymmetry in segment width corresponds to an asymmetry in the amount of spatial territory covered, further supporting a sensory-motor topographic map for a sucker in the ANC.

### Comparative analysis

We carried out a comparative analysis to clarify the relationship between ANC segmentation and function by examining the organization of the ANC of the longfin inshore squid, *Doryteuthis pealeii*. Squids, which diverged from octopuses 270 million years ago, have two tentacles in addition to eight sucker-lined arms^20^ (Fig 4a, e, i). Like octopuses, these limbs are muscular hydrostats, yet they differ in gross morphology and function^21,22^. Where *O. bimaculoides* is benthic, with arms suited for exploration and movement on the seafloor, *D. pealeii* is a pelagic animal and primarily uses its arms and tentacles for prey capture in open water^7^. The tentacle stalks, which are devoid of suckers, rapidly elongate to grab prey with tentacle clubs, the sucker-rich ends of the tentacles. The arms enclose the prey in a coordinated movement as the prey is brought back to the mouth. Morphological differences between squid arm and tentacle muscle composition underlying this functional difference have been described^22,23^.

**Figure 4:**
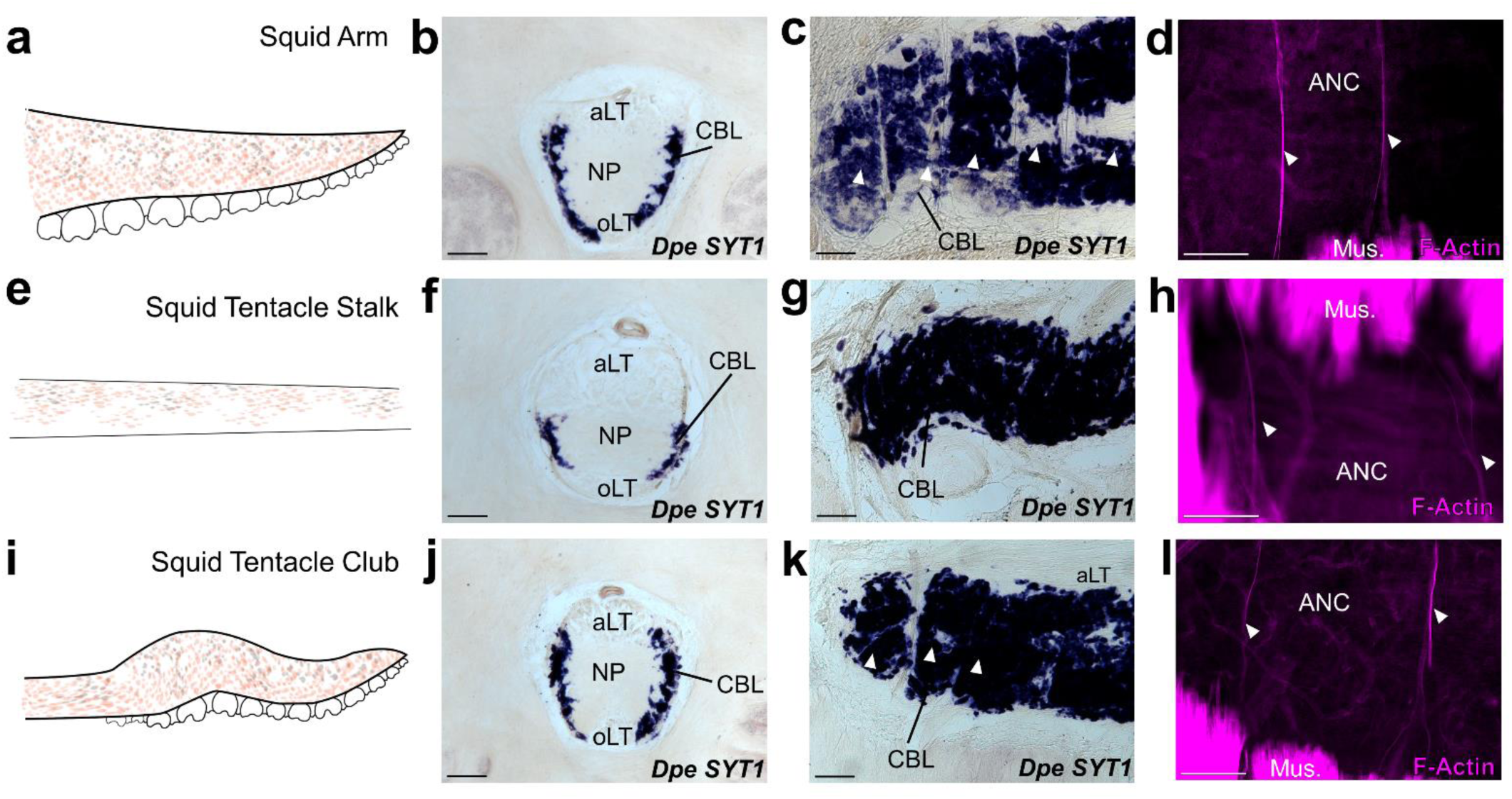
Segmentation is a shared feature of flexible, sucker-laden cephalopod appendages. a-d. The ANC of the *D. pealeii* arm is segmented. **(a)** Cartoon depiction of *D. pealeii* arm, which has suckers. **(b)** ISH for *Dpe SYT1* of an ANC transverse section demonstrating the CBL. **(c)** ISH for *Dpe SYT1* of an arm longitudinal section demonstrates CBL segmentation. **(d)** F-actin (magenta) marks vasculature in the arm ANC septa. **e-h** The ANC of the tentacle stalk lacks CBL segmentation. **(e)** Cartoon depiction of *D. pealeii* tentacle stalk which is devoid of suckers. **(f)** Transverse ISH section for *Dpe SYT1* demonstrates a clear, although reduced, tentacle CBL. **(g)** *Dpe SYT1* ISH of a longitudinal ANC section does not show CBL segmentation in the tentacle stalk. **(h)** F-actin (magenta) demonstrates a regularly arranged vasculature in the tentacle stalk, but these blood vessels do not lie within the CBL. **i-l,** The ANC in the tentacle club has segments. **(i)** Cartoon depiction of *D. pealeii* tentacle club, which is located at the end of the stalk and has suckers. **(j)** Transverse ISH section for *Dpe SYT1* in the tentacle club ANC demonstrates the CBL. **(k)** Longitudinal *Dpe SYT1* section of the tentacle club CBL shows segmentation. **(l)** F-actin (magenta) identifies regularly in the tentacle club septa. White arrowheads in **(c, k)** point to CBL segments. Those in **(d, h, l)** indicate phalloidin-labeled blood vessels. aLT, aboral longitudinal tract; oLT, oral longitudinal tract. Scale bars: 100 μm.

In both the arms, tentacle stalks and clubs of *D. pealeii*, there is an ANC mediating sensorimotor control (Fig 4b, f, j). The ANC in the squid has a notably different arrangement from that of octopus, however. The CBL is restricted to the ANC lateral edges, and there are large longitudinally running fiber-tracts both orally and aborally (Fig 4b, f, j). The CBL thins along the tentacle stalk but expands in the sucker-rich tentacle club (Fig 4f, j). We used gene expression and phalloidin staining to look for segmentation in the squid CBL. The CBL in the arm of the squid is segmented, with approximately 4.5 segments per sucker (Fig 4c, d). As in *O. bimaculoides*, F-actin is localized to the ANC septa (SFig 8a). In the tentacle stalk, CBL segments as identified by *SYT1* gene expression are at best indistinct, and although F-actin marks regularly spaced vasculature, these are not in the CBL (Fig 4g, h, SFig 8b). In the sucker-rich tentacle club, by contrast, CBL segments marked by *SYT1* and F-actin within the CBL are readily identified (Fig 4k, l, SFig 8c). This comparative anatomy suggests a strong link between ANC segmentation and flexible cephalopod arms lined with suckers.

## Discussion

The arm is a highly redundant structure, with both suckers and the pattern of brachial musculature repeating down its length^1,2^. A parcellation of the ANC into segments is a natural way to relegate motor control of a continuous limb with high redundancy. In fact, many computational models and soft robots inspired by octopus arms divide the arm into repeated segments in their construction or control algorithms^24–27^. Our results, however, demonstrate additional complexities. For the brachial musculature, different sets of nerves exit from adjoining septa, while still providing coverage of the musculature. Instead of a single segment representing the full cylinder of brachial musculature, as is the case in many models, adjoining segments coordinate in parsing control for the muscles. This circuit structure could more effectively smooth continuous movements along the length of the arm, as well as isolate contractions to localized territories in the brachial musculature.

The ANC contains oscillating enlargements corresponding to the location of the suckers. The neuronal segments described here further subdivide these larger swellings into multiple modules. Classic studies have proposed that much of the sensorimotor processing in suckers occurs within the epithelium or in the sucker ganglion, and that by the time the information reaches the ANC, it has been highly downsampled^28,29^. Our investigation of nerve fiber connections illustrates that, at the least, spatial information is preserved within the ANC, creating a suckerotopy. Combined with the projections that interconnect suckers, such a neural architecture would support many behaviors seen in suckers, from passing objects along suckers to anisotropic detection of chemical cues^5,30–32^. In particular, shape deformation of suckers has been thought to underlie texture discrimination^33^. This spatial mapping could be the basis for ring attractor computations by the axial nerve cord to guide sucker movements in response to sensory cues. Such a mechanism has been described for spatial navigation in the central complex of *Drosophila*^34^, and recently it has been suggested in biophysical models of directed movements by arms^35^.

Molluscs are a diverse phylum, and cephalopods in particular exhibit many derived morphologies. Our comparative analysis emphasizes that this segmentation is a derived feature in flexible, sucker-laden arms. Segments are present in the arm and tentacle club in *D. pealeii,* yet they are indistinct in the stalk which is devoid of suckers. Interestingly, there are fewer segments per sucker in the arm of *D. pealeii* compared with that of *O. bimaculoides.* Differences in ecological niche and behavioral repertoire, including prey hunting strategies, could drive this variation^7,32^.

There has long been a vigorous discussion in the evolutionary and developmental biology fields about whether segmentation is an ancestral feature of bilateral animals^36–41^. In particular, segmentation in molluscs has been proposed for basally branching polyplacophorans (chitons) and monoplacophorans^36^. This has, however, been hotly contested and at the least has not been proposed to encompass the nervous system^37,39–42^. A second view of segmentation is that it represents an adaptation useful for generating vermiform movements^41,43^. Our findings provide a clear example of the second view, that of derived segmentation linked to the control of the motor patterns characteristic of soft-bodied cephalopod (coleoid) arms. This segmentation shares many similarities to segmentation described in other systems: multiple tissue types, reiterated patterning along a longitudinal axis, and utilization for sensorimotor control circuitry^13,14,19,41^. At its core, however, the coleoid segmentation appears to reflect one role, that of setting up recurring neuronal modules dedicated to the control of muscles and suckers which are themselves reiterated down the length of the arms.

## Materials and Methods

### Animals

Wild caught adult *Octopus bimaculoides* were purchased from Aquatic Research Consultants, a field-collection venture operated by Dr. Chuck Winkler, San Pedro, CA. Animals were individually housed in 20-gallon saltwater tanks equipped with a carbon particle filter, aquarium bubbler, and an UV light.

Animals were fed daily with fiddler crabs. The artificial seawater (ASW) was prepared in deionized water from pharmaceutically pure sea salt (33g/liter; Tropic Marin “classic”, Wartenberg, Germany). Adults were deeply anesthetized in 4% ethanol/ASW (EtOH/ASW, n = 17) and trans-orbitally perfused with 4% paraformaldehyde/phosphate-buffered saline solution (PFA/PBS, pH 7.4) delivered via a peristaltic pump (Cole Parmer Masterflex) through a 21½ gauge needle (Becton Dickinson). The left and right white bodies, a hematopoietic tissue, were targeted alternately and iteratively throughout the procedure. Arms were dissected and stored in 4% PFA/PBS overnight at 4°C. Specimens were then either cut into 2-4 cm pieces for immediate processing or stored in PBS at 4°C.

Wild caught adult *Doryteuthis pealeii* (n = 3) were obtained from the Marine Biological Laboratory, Woods Hole, MA. Animals were kept in circulating, filtered containment tanks for several days before being deeply anesthetized in 7.5% MgCl2/ASW and dissected. Arm crowns were immersion fixed in 4% PFA/PBS. Arms and tentacles were dissected and cut into 2-4 cm pieces for further processing.

These cephalopod experiments were performed in compliance with the EU Directive 2010/63/EU guidelines on cephalopod use, the University of Chicago Animal Resources Center and the MBL and UChicago Institutional Animal Care and Use Committees^44,45^.

### Tissue Processing

For sections, arm explants were incubated in 30% sucrose/4%PFA/PBS at 4°C for three to five days until saturated, rinsed with 30% sucrose/PBS, and infiltrated with 10% gelatin/30% sucrose/PBS for 1 hour at 50°C. Tissue was embedded in 10% gelatin/30% sucrose/PBS and post-fixed in 30% sucrose/4%PFA/PBS before storage in -80°C. Gelatin-embedded serial arm sections were cut at 28-50-µm thickness on a freezing microtome (Leica SM2000R) and collected in diethyl pyrocarbonate (DEPC)-treated PBS. Sections were mounted and dried on charged, hydrophilic glass slides (TruBond380, Newcomer Supply, Middleton, WI). Slides were stored at -80°C until further processing.

### Immunohistochemistry

Neuronal processes in octopus arm were labeled with mouse monoclonal antibody 6–11B-1 (1:500 dilution of ascites fluid; Sigma-Aldrich, Cat#: T6793) and mouse monoclonal antibody SMI-31 (1:500 dilution; BioLegend, Cat# 801601). Clone 6–11B-1 was isolated following immunization with sea urchin sperm flagella protein preparations^46^. It recognizes an acetylated α-tubulin (acTUBA) epitope found broadly but not universally across microtubules and has been extensively employed to identify axon tracts in vertebrate and invertebrate nervous systems^47–50^. Clone SMI-31 reacts with a phosphorylated epitope of neurofilament in mammals and neurofilament 220 in squid^51–53^ .The secondary Alexa Fluor® 488 AffiniPure Goat Anti-Mouse IgG (Cat#: 115-545-003) and Cy™3 AffiniPure Donkey Anti-Mouse IgG (Cat#: 715-165-151) were purchased from Jackson ImmunoResearch (West Grove, PA) and employed at 1:500 dilutions.

For section immunohistochemistry, slides were rinsed 3×10 minute in PBS containing 1% tween 20 (PBST) and incubated for 30 minutes in a proteinase K solution (Sigma-Aldrich, Cat#: 03115828001; 19.4 μg proteinase K per milliliter of PBST). Slides were post-fixed with 4% PFA, washed 3×30 minutes in PBST, blocked in 10% goat serum/PBST (Fisher Scientific, Cat#: 16210072) for 1 hour and incubated in primary antibody diluted in 1% goat serum/PBST for four days at 4°C. After 3×30 minute PBST washes, sections were incubated for two days at 4°C with secondary antibody along with 0.01 mg/ml 4′-6-diamidino-2-phenylindole (DAPI; Sigma-Aldrich, Cat#: D9542) to label cell nuclei fluorescently and either 0.2 µl/ml phalloidin-iFluor 594 (Abcam, Cat#: ab176757) or phalloidin-iFluor 488 (Abcam, Cat#: ab176753) to label F-actin. Slides were rinsed and washed 3×5 min with PBST and cover slipped with Fluoromount G (Southern Biotech, Birmingham, AL).

For whole mount immunohistochemistry, arm slice explants were prepared: dissected axial nerve cord, 0.5 cm – 1 cm transverse slices of arm, longitudinally bisected slices, and horizontally bisected slices.

Explants were washed 3×15 minutes in PBST, dehydrated in a graded methanol series (25, 50, 75% in PBST, each for 10 minutes), rinsed twice is 100% methanol for 10 minutes, and stored overnight at -20°C. The next day, explants were rehydrated in a graded methanol series (75%, 50%, 25% in PBST, each for 5 minutes) and rinsed twice in PBST for 5 minutes. Then, tissue was incubated for 60 minutes at 37°C in a proteinase K solution (19.4 μg proteinase K per milliliter of PBST), post fixed for 15 minutes in 4% PFA/PBS, washed 3×15 min and 1×60 min in PBST, and blocked for an hour in 10% goat serum/PBST. Slices were then incubated in primary antibody for 7 days at 37°C. Following primary incubation, tissue was rinsed 1x, 3×15 minutes, 5×60 minutes and overnight in PBST. Tissue was transferred to secondary antibody for seven days at 4C. Next, the tissue was rinsed with PBST 1x, 3x15, 2×60 minutes, washed with PBS 3×15 minutes, and post fixed for four days at 4°C. Explants were then rinsed 3×15 min in PBS and cleared in a modified version of FRUIT (incubations in 35, 40, 60, 80, 100% FRUIT, each for 24 hours)^54^. Slices were stored in 100% FRUIT at 4°C until imaged.

### Picrosirius Red

Collagen in squid and octopus samples was examined with Picrosirius Red (Abcam, ab150681). Gelatin was melted off at 72°C for 4-6 hours before slides were dried and processed at room temperature. Slides were rinsed 3 times with deionized water, before being rehydrated for 1 minute. Sections were incubated in Picrosirius Red for 5-10 minutes. Slides were de-stained with two rinses of acetic acid wash, before being dehydrated in 100% EtOH and mounted with Eukitt (Sigma Aldrich).

### Hematoxylin and Eosin Staining

Gelatin was melted off sectioned tissue at 72°C for 4-6 hours before slides were dried and processed at room temperature. Slides were rehydrated in deionized water and incubated in Mayer’s Hematoxylin (HAE-IFU, ScyTek) for 1 minute, before rapid rinsing in water to prevent overstaining. Tissue was briefly incubated in a bluing solution (HAE-IFU) for 15 seconds. The tissue was then dehydrated in an ethanol series (70%, 95% and 100%) before being moved into Eosin Y (HAE-IFU) for 1 minute. Slides were rinsed in 100% EtOH, cleared in Histoclear (National Diagnostics, Atlanta, GA) for 1 minute and mounted with Eukitt.

### cDNA Synthesis and Cloning

Dissected octopus central brain tissue and arm tissue was flash-frozen on dry ice and stored at -80°C until RNA extraction. Tissue was homogenized with a micropestle, and RNA was extracted with Trizol Reagent (Invitrogen) and phasemaker tubes (Invitrogen) following manufacturer’s instructions. RNA was stored at -80°C in RNase-free water (Sigma-Aldrich) until used for cDNA synthesis with SuperScript III 1st-strand cDNA kit (Invitrogen) following manufacturer instructions. cDNA was diluted in RNase-free water and stored at -20°C.

PCR primers were designed with MacVector software (version 12.6.0) or PrimerBlast from NCBI (Table 1). PCR reactions were conducted using the T100 thermocycler from BioRad. Reaction solutions were incubated at 95°C for 5 minutes before undergoing 35-40 rounds of amplification cycles: 95°C for 30 seconds, 52-57°C for 45 seconds, and 72°C for 1 minute. A final elongation step was performed at 72°C for 10 minutes. The sequence for *Dpe SYT1* was synthesized by Twist Bioscience (South San Francisco, CA; Table 1).

**Table 1.**
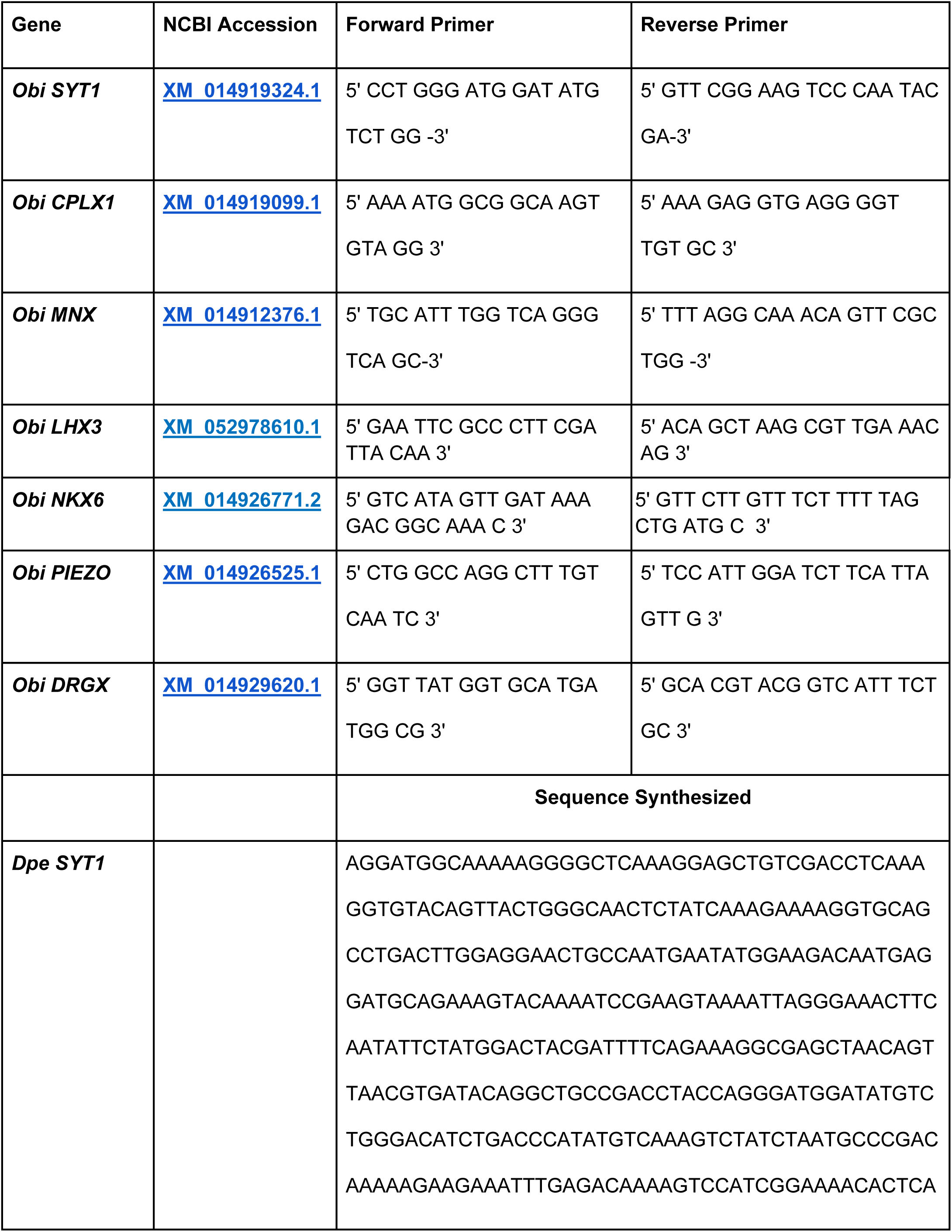
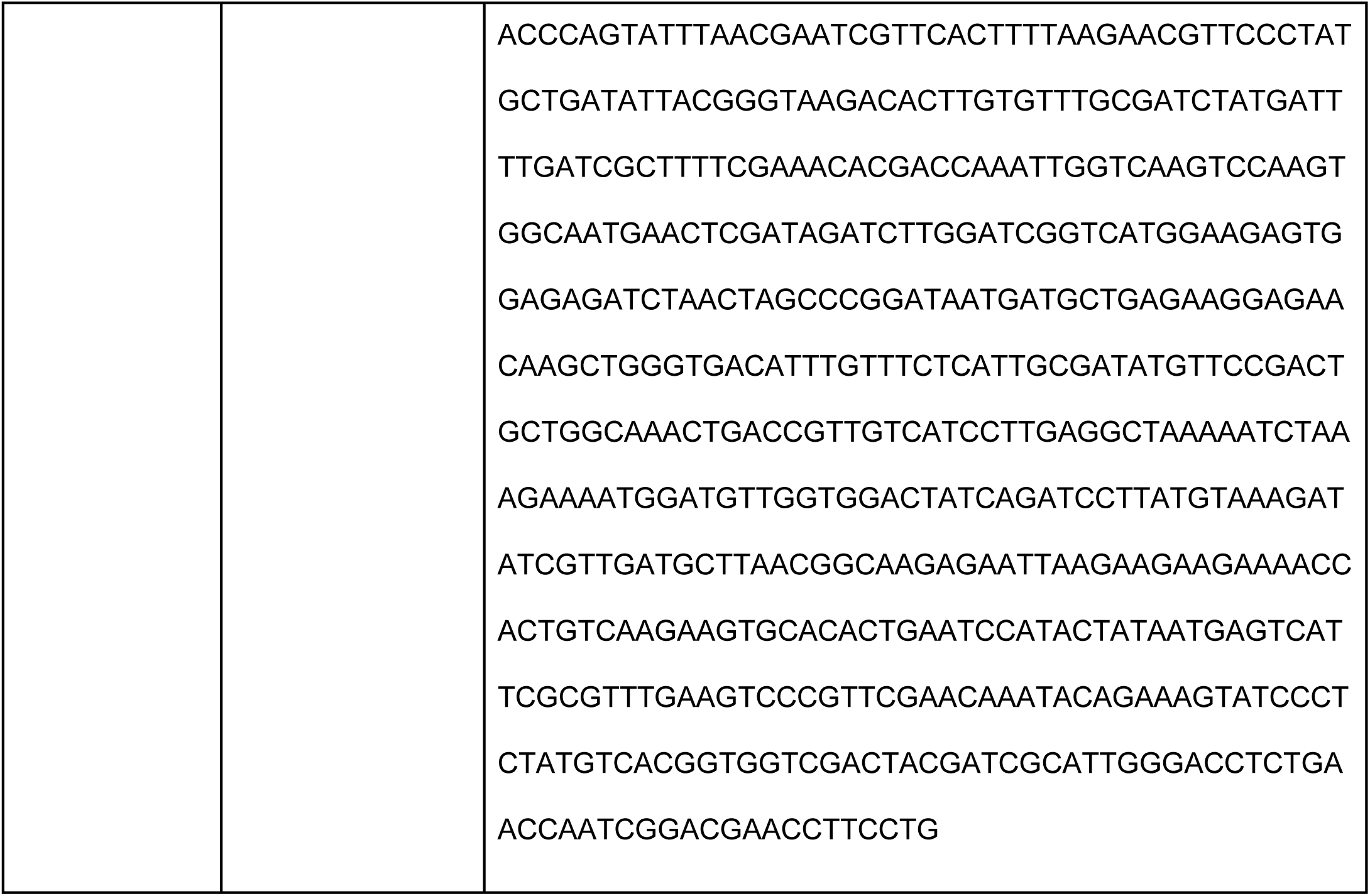

PCR products and Twist sequence were studied by gel electrophoresis to confirm that the sizes of products were as expected. PCR reactions were ligated into a pGEM T-Easy plasmid (Promega). Closed inserts were Sanger sequenced by the UChicago Comprehensive Cancer Center DNA Sequencing Facility. Plasmids were linearized by SacII (New England Biolabs, Cat#: 50812058) or SpeI (New England Biolabs, Cat#: 50811989) restriction enzyme digestion to generate antisense templates. Following phenol-chloroform extraction of the template, antisense digoxigenin (DIG)-labeled riboprobes (Sigma-Aldrich, Cat#: 11277073910) were transcribed with SP6 or T7 RNA polymerase (New England Biolabs). After transcription, residual template was digested with RNase-free DNase I (Sigma-Aldrich) at 37°C for 45 minutes. Riboprobes were ethanol-precipitated and stored in 100μl of DEPC-H_2_O at -20°C until use.

### In situ hybridization

Slides of sectioned tissue were equilibrated to room temperature and post-fixed in mailers for 15 minutes in 4% PFA/PBS, washed 3x 15 minutes in DEPC-PBS and incubated at 37°C for 15 minutes in proteinase K solution (Sigma-Aldrich; 1.45 μg proteinase K per milliliter of 100 mM Tris-HCl [pH 8.0], 50 mM EDTA [pH 8.0]). Slides were post-fixed for 15 minutes in 4% PFA/PBS, washed 3x 15 minutes in DEPC-PBS, and acclimated to 72C for one hour in hybridization solution (50% formamide, 5x SSC, 1% SDS, 200 μg/ml heparin, 500 μg /ml yeast RNA). Slides were transferred to mailers with 1-2 mg antisense riboprobe in 15 mL hybridization solution and incubated overnight at 72°C. The next day, slides were treated 3x 45 minutes in preheated Solution X (50% formamide, 5x SSC, 1% SDS) at 72°C. Slides were washed 3x 15 minutes in room temperature TBST (Tris-buffered saline with 1% Tween 20) and blocked at room temperature for one hour in 10% DIG buffer (Roche) in TBST.

Anti-DIG Fab fragments conjugated to alkaline phosphatase (Sigma-Aldrich, Cat#: 11093274910) were preabsorbed with octopus embryo powder in 1% DIG buffer in TBST for at least one hour. Slides were then incubated on a rocker overnight at 4°C in preabsorbed antibody diluted to a final concentration of 1:5000 in 10% DIG buffer in TBST.

The next day, slides were rinsed once, washed 3x 15 minutes, then 2x 1 hour in TBST. Slides were washed for 10 minutes in freshly prepared NTMT (100 mM Tris-HCl [pH 9.5], 100 mM NaCl, 50 mM MgCl2, 1% Tween 20). For the color reaction, slides were incubated in nitro blue tetrazolium (NBT, 100mg/mL in 70% dimethyl formamide/30% DEPC-H20, Gold Biotechnology, St. Louis, MO) and 5-bromo-4-chloro-3-indolyl phosphate (BCIP, 50mg/mL in 100% dimethyl formamide, Gold Biotechnology, St. Louis, MO) in NTMT. Color development proceeded at room temperature and was monitored for a maximum of 5 days. When reaction was complete, slides were washed in TBST overnight and dehydrated through a series of ethanol washes into Histoclear, and coverslipped with Eukitt (Sigma-Aldrich).

### Tracing

Adult *O. bimaculoides* (n = 13) were anesthetized in 2% EtOH/ASW, and the distal portion of a single arm was amputated with a razor blade. The amputated arm was placed in ASW, and the animal was either returned to a bucket of ASW to recover or euthanized in 4% EtOH/ASW and perfused.

The amputated arm was cut into 0.5cm-1cm slices. Each slice (n = 83) was injected with CF® 488A Dye Dextran (Biotium, Cat#80110; 2% solution, 10kD) using a 25 or 32 gauge Hamilton Syringe (Hamilton, model#: 7000.5), rinsed in filtered seawater, and incubated at room temperature in 221 media with NuSerum (36% Leibovitz L15-Media, 36% filtered seawater, 18% deionized water, 1% pen-strep, 10% NuSerum). After three hours, the slices were rinsed with filtered seawater and fixed in 4% PFA/PBS. Slices were prepared for gelatin embedding and sectioning as described above, sectioned at 50um, and counterstained with phalloidin-iFluor 594 (0.2 µl/ml in 1%PBST) and DAPI (0.01 mg/ml in 1% PBST) before imaging.

### Vasculature Injections

Lab-raised 6-week-old *O. bimaculoides* (n = 5) were anesthetized in 2% EtOH/Filtered ASW. A small incision was made above the right orbital socket to provide access for the pulled glass electrode to enter the white body. Using a picopump, 3-5 μL of a heavy-weight dextran (Invitrogen, Cat#D1818, 70,000 MW, lysine fixable) was injected into each juvenile octopus and allowed to circulate throughout the body for 5 minutes. Animals were then transferred to a 4% EtOH/Filtered ASW for five minutes before immersion fixation in 4% PFA/PBS overnight at 4°C. Injected animals were stored in PBS at 4°C before further processing for imaging or immunostaining.

### Imaging

The immunolabeled and tracing tissue was studied with a Zeiss Axioskop 50 upright microscope and a Leica MZ FLIII stereomicroscope, both outfitted with the Zeiss AxioCam digital camera and AxioVision 4.5 software system. Selected sections were also studied on a Leica SP5 Tandem Scanner Spectral 2-photon confocal microscope (Leica Microsystems, Inc., Buffalo Grove, IL) or scanned with an Olympus VS200 Research Slide Scanner (Olympus / Evident, Center Valley, PA) with a Hamamatsu ORca-Fusion camera (Hamamatsu Photonics, Skokie, IL). Whole mounts were imaged on a LaVision BioTec UltraMicroscope II (Miltenyi Biotec, Bergish Gladbach, Germany) run by ImSpector Pro v. 7_124 software (LaVision BioTec, Bielefeld, Germany). Collected images were corrected for contrast and brightness and false-colored FIJI (version 2.1.0/1.53c; National Institutes of Health (NIH)). Whole mount reconstructions were visualized with Arivis Vision4D software v. 3.1 (arivis AG, Rostok, Germany).

### Proximal-distal analysis

To examine the segmentation pattern down the proximal-distal axis, two series were created: 3 evenly spaced arm explants from an arm on the right side of one animal sectioned longitudinally and stained with acTUBA, and 3 evenly spaced explants from an arm on the left side of a second animal sectioned longitudinally and stained with H&E. For cross sectional area, an arm explant was sectioned horizontally and stained with acTUBA. For segment counts on *D. pealeii*, counts were performed on sections of arm explant stained with ISH for *Dpe SYT1*.

#### Segment counts

The number of segments were counted along a length of six suckers on the anterior and posterior side of the ANC. The counts for the anterior and posterior sides were averaged together, then divided by six to determine segments/sucker.

#### Segment Width

Four segments across six suckers on both the anterior and posterior side of the ANC were selected for width measurements and were tagged with either the external side of the ANC (ExA) or internal side of the ANC (InA) (SFig 7a). Width of each segment along the proximal-distal axis was measured manually using the line tool in FIJI. Measurements on the anterior and posterior side were averaged together to get averages for ExA and InA. Total averages were found by additionally averaging ExS and InS.

#### Sucker Width

For six suckers in each explant, the width of the acetabulum was measured manually at the widest point using the line tool in FIJI.

#### Cell Density

For each of the segments selected for the width measurements, a rectangle spanning the width of the segment was taken. The number of nuclei, visualized by DAPI, within the rectangle were counted using the analyze particle function in FIJI. The area of the rectangle was also found in FIJI. Density was calculated as the number of nuclei/area. Averages for ExA and InA pooled across both the anterior and posterior side of the ANC as for segment width calculations.

#### Cross sectional area

Across three suckers, three segments on ExA and three segments on InA were selected for cross sectional area measurements. Visibility of each segment across the oral to aboral extent of the ANC were ensured. The width, defined as the length in the proximal-distal axis, and height, defined as the length perpendicular to the proximal-distal axis, were manually measured in FIJI. Cross sectional area was calculated as width x height.

### Nerve analysis

Large scale nerve analysis was done by segmenting whole mount tissue, immunostained for nerve markers, acTUBA and SMI-31. To examine brachial nerves, two whole mounts of slices stained with acTUBA and one slice whole mount stained with SMI-31 were imaged transversely. To examine the oral nerves, two whole mounts labeled for acTUBA were imaged, one transversely and one longitudinally, and one whole mount with SMI-31 was imaged transversely. Nerve fibers going to the sucker and to the brachial musculature were traced using Simple Neurite Tracer (SNT) in FIJI^55^.

#### Oral nerve analysis

The traces for the oral nerves were tagged as corresponding to ExA or InA and the 3D spatial coordinates outlining the traces were found using the SNT API (SFig 7a, g)^55^. The coordinates of tips of the nerve were translated such that the center of all the tips was set as the origin. Subsequently, each tip coordinate was normalized such that the vector from the origin to the tip has a magnitude of 1. The spatial distribution of tips corresponding to the ExS and the InS were visualized using Matplotlib.

#### Aboral and central nerve analysis

The traces for the brachial nerves were segregated into the central nerves targeting the oral brachial musculature and the aboral nerves targeting the aboral brachial musculature (SFig 4a). The branching pattern of the nerves were characterized using the SNT Sholl analysis with 10 μm spacing (SFig 4b)^56^. The profile was then fit with a 5-degree polynomial. The 3D spatial coordinates, root, and tips of the nerves were found using the SNT API^55^. The average trajectory of the nerve was computed by creating a vector from the root of the nerve to the center of the process, and the distribution of the tips in the aboral-oral and proximal-distal plane were visualized using Matplotlib (SFig 4c, 5a). Both the Sholl plots and average trajectory were normalized such that the length of the longest nerve, the nerve that reaches the skin, was set 1. Relation to the septa were determined using the oral roots as additional markers and used to color code the nerves.

### Statistics

Unless otherwise stated, data are mean +/- standard error of the mean (sem). Using MATLAB, a two-way ANOVA followed by Tukey’s post hoc test was conducted to test for significance along the proximal-distal axis and between the ExS and InS. Data were considered significant if p < 0.05.

## Data availability

Data are available from the corresponding author upon request.

## Code availability

Custom code used to analyze proximal-distal series and nerve fiber patterns are available from the corresponding author upon request.

## Acknowledgements

We thank Dr. Chuck Winkler of Aquatic Research Consultants for providing us with octopuses and Drs. Caroline Albertin and Thea Rogers for the *D. pealeii* samples. Imaging was performed at the University of Chicago Integrated Light Microscopy Core (RRID: SCR_019197). We extend special thanks to Dr. Christine Labno and Mr. Khalil Rodriguez for their invaluable assistance. We also thank Ms. Caroline Miller for assistance with whole mount image processing, and Ms. Amelia Cheng and Mr. Jan Kasal for assistance with tracer injections and tissue processing. This work was supported by the NIH Developmental Biology Training Program (HD055164) and the NSF GRFP (NGS), and by the NIH UF1NS115817 award (CWR).

## Supplemental

**Supplemental Figure 1:**
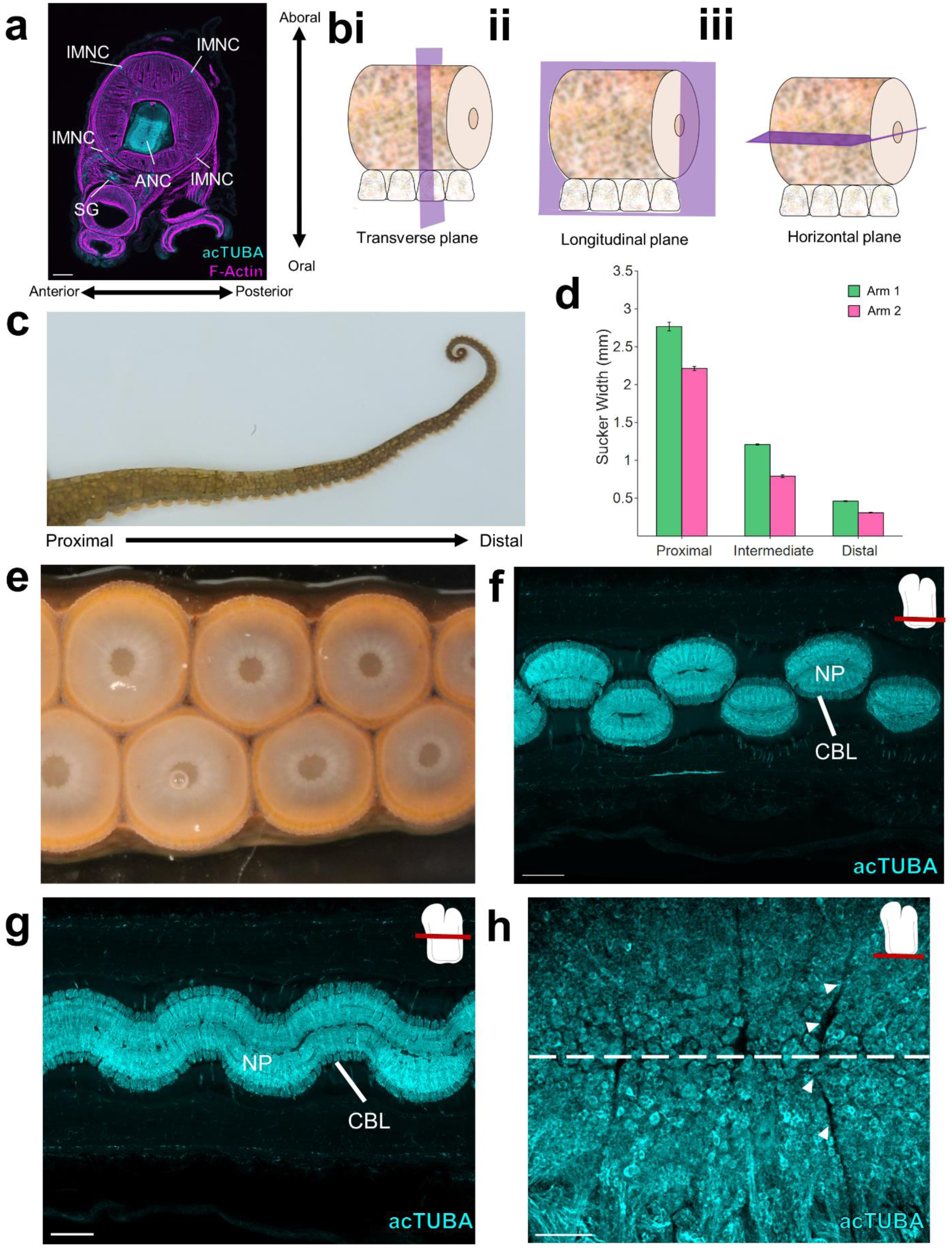
Arm anatomy overview. **a,** Transverse section of the arm stained with acTUBA (cyan) and phalloidin (F-Actin, magenta). The main components of the arm nervous system are highlighted. ANC: Axial nerve cord, IMNC: intramuscular nerve cord, SG: sucker ganglion. Scale bar: 500 μm. **b,** The three perpendicular planes of sectioning through the arm. **i,** The transverse plane is perpendicular to the long axis of the arm. **ii,** The longitudinal plane is parallel to the long axis of the arm. **iii**, the horizontal plane is parallel to the long axis of the arm and to the suckers. **c,** Photograph of the arm of *O. bimaculoides.* From proximal to distal, the girth of the arm decreases. **d,** Sucker width decreases from proximal to distal. Arm 1 (green) and Arm 2 (pink). n = 6 suckers per condition, error bars +/- sem. **e,** An en face photograph of the suckers shows that the suckers are arranged in two offset rows. **f,** Horizontal section of the ANC through the sucker territory labeled with acTUBA (cyan). The ANC oscillates to the side overlying the sucker. Scale bar: 500 μm. **g,** Horizontal section of the ANC through the brachial territory labeled with acTUBA (cyan). Scale bar: 500 μm. **h,** Maximum projection of a horizontal slice whole mount immunolabeled with acTUBA (cyan) through the oral CBL. Segments extend across the midline (denoted with the dashed line). Scale bar: 100 μm.

**Supplemental Figure 2:**
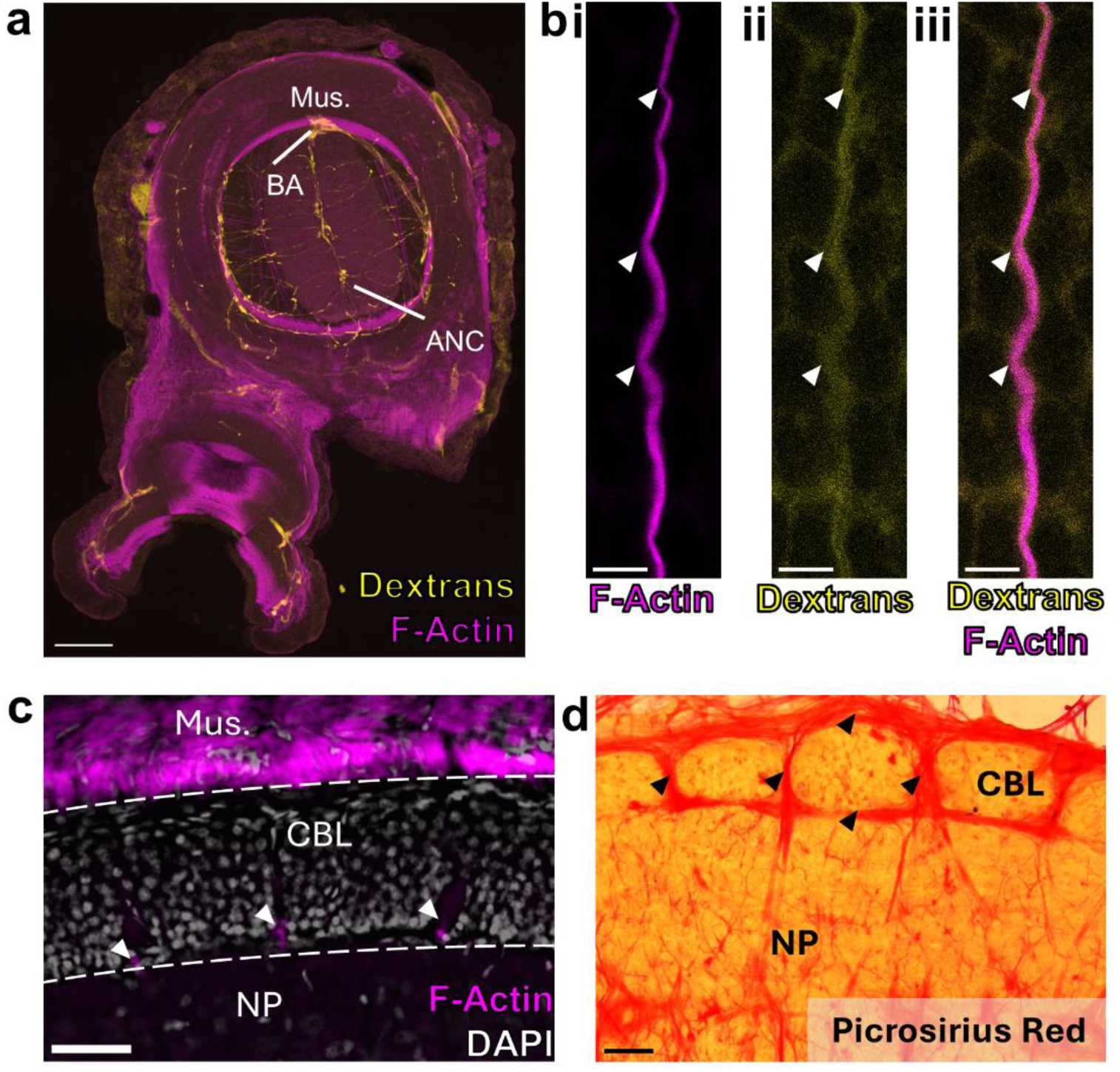
Septal localization of vasculature and collagen. **a,** Transverse section of arm with dextran labeling (yellow) of the vasculature. F-actin and dextran delivery clearly identify the large brachial artery (BA). Scale bar: 100μm. **b,** Maximum projection of a single blood vessel. F-actin (magenta, **i**) and dextrans (yellow, **ii**) colocalize (**iii**). Scale bar: 5 μm. **c.** Horizontal section through the ANC labeled with F-actin (magenta) and DAPI (gray). The vessels are located in the CBL at the interface between the CBL and the neuropil (NP). Scale bar: 50 μm. **d,** Horizontal section through the ANC labeled with Picrosirius Red. Collagen, labeled in red, wraps around the CBL segments and extends within the full length of the septa. Scale bar: 50 μm.

**Supplemental Figure 3:**
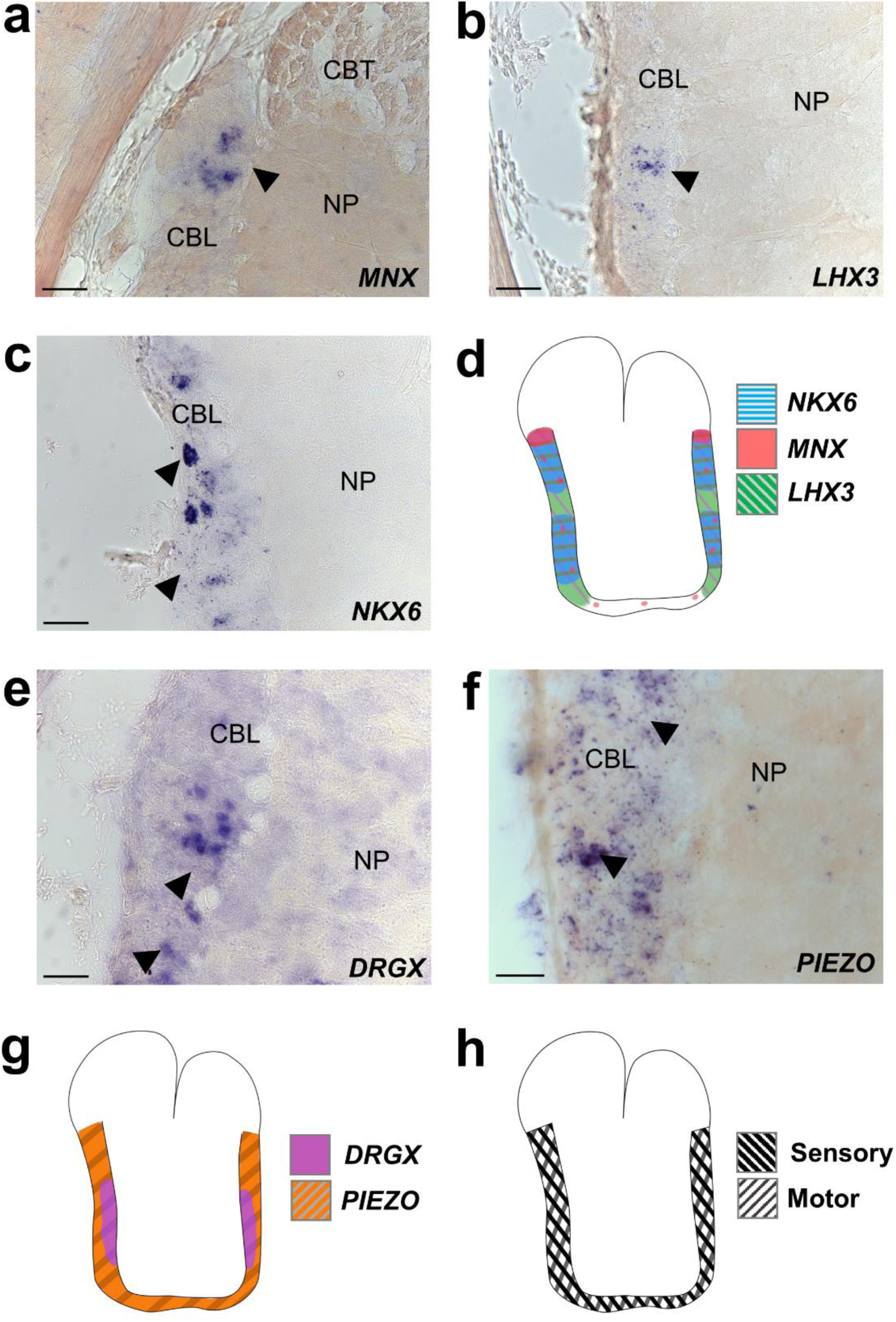
Markers of sensory and motor neurons label overlapping territories in the ANC. **a-d,** ISH for motor neuron markers in ANC transverse sections. **(a)** *MNX*, **(b)** *LHX3*, **(c)** *NKX6*. **(d)** Cartoon summary of motor neuron marker distributions. Expression is extensive in the lateral walls of the CBL. **e-g,** Expression of sensory neuron markers in ANC transverse sections. **(e)** DRGX, **(f)** *PIEZO.* **(g)** Cartoon summary of sensory neuron marker distribution. Expression is also strong in the CBL lateral walls. **h,** Diagram of overlapping sensory and motor neuron territories in the CBL. Scale bars: 50 μm.

**Supplemental Figure 4:**
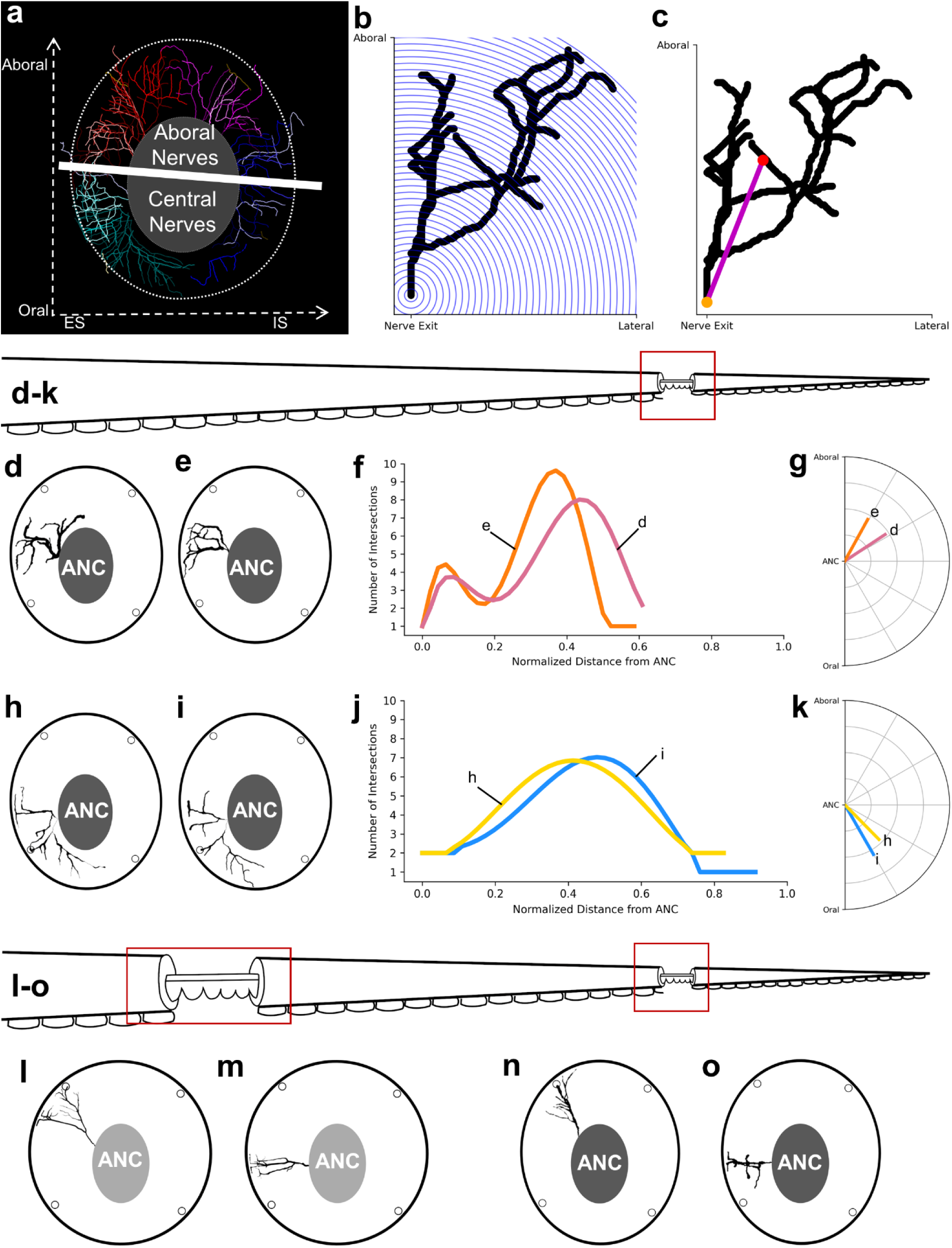
Nerve fiber analysis. **a,** Brachial nerves segmented out of transverse whole mount slice immunolabeled with acTUBA. Brachial nerve fibers can be split into aboral nerves and central nerves based on ANC exit location. **b,** Diagram depicting Sholl analysis on an example nerve. Branching is characterized as the number of intersections between the nerve fiber and a series of spheres centered at the nerve exit, with an increasing radius of 10 μm. Three-dimensional data were analyzed and are captured here in two-dimensions. **c,** Schematic of the average trajectory, which is a vector created from the nerve exit point to the center of the processes. **d-k**, Within the same slice, similarities in nerve fibers are found separated by more than one segment. **d, e,** Transverse diagrams with two aboral nerves that have similar morphologies. These nerves also have **(f)** similar Sholl profiles and **(g)** average trajectories. **h, i,** Transverse diagrams with two central nerves that have similar morphologies. These nerves also have **(j)** similar Sholl profiles and **(k)** average trajectories. **l-o,** Similarities in nerve fiber morphologies can be found when comparing proximal and distal slices. An aboral nerve from a proximal slice in **(l)** is similar to a selected aboral nerve from a distal slice in **(n).** A central nerve from a proximal slice in **(m)** is similar to a selected central nerve from a distal slice in **(o)**.

**Supplemental Figure 5:**
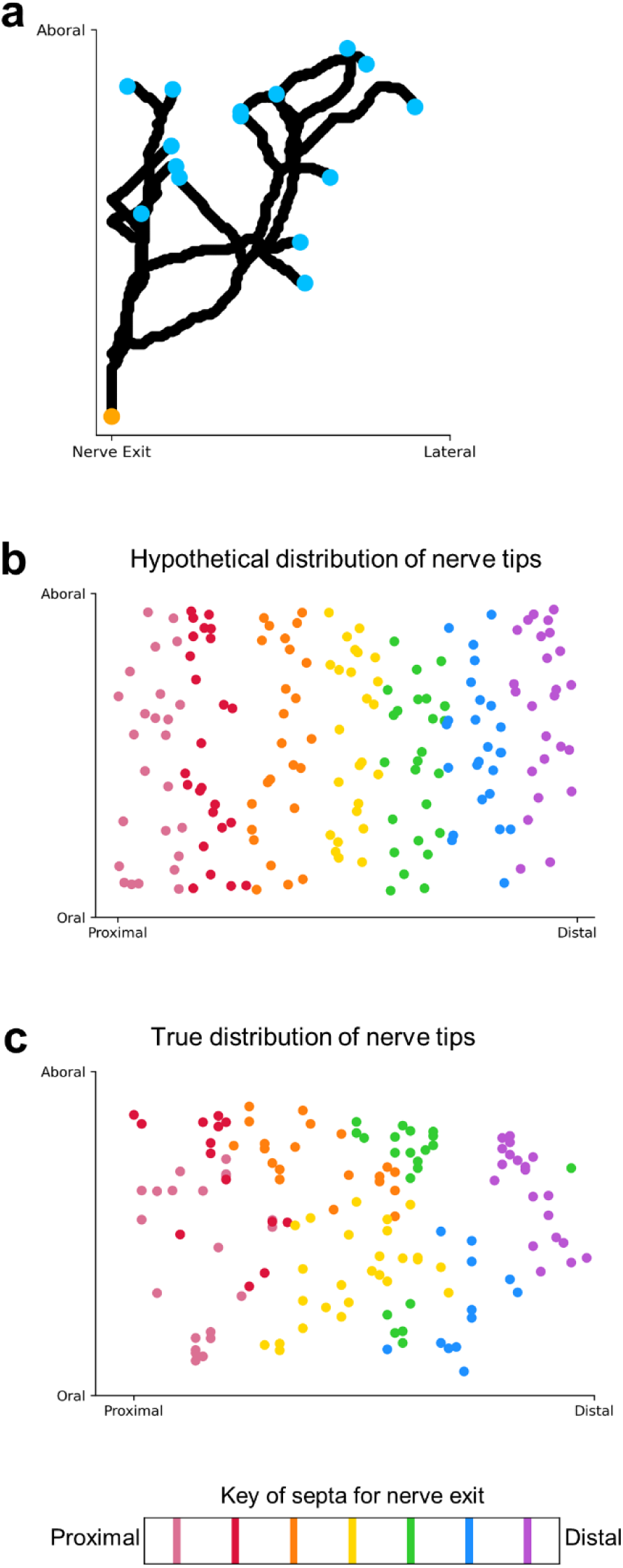
Distribution of brachial nerve tips. **a,** Example brachial nerve with nerve tips labeled. **b,** Hypothetical distribution of nerve tips across the proximal-distal axis, colored by ANC exit point. Nerves exiting from one septum fully cover the aboral-oral extent of the brachial musculature and do not intercalate with nerves exiting from other septa. **c,** True distribution of nerve tips across the proximal-distal axis, colored by ANC exit point. Only nerve fibers from multiple septa added together cover the aboral-oral extent of the brachial musculature.

**Supplemental Figure 6:**
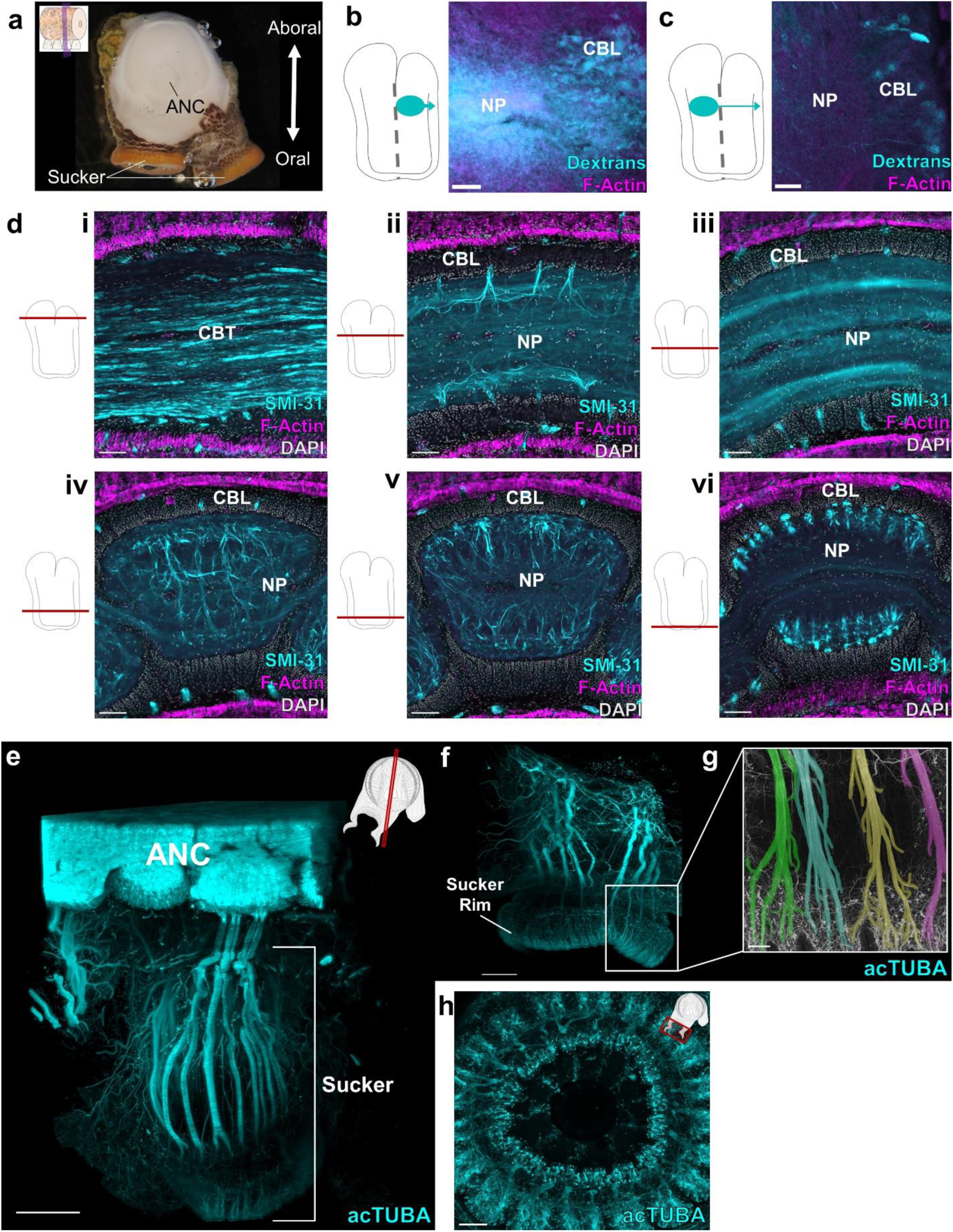
Neuropil organization. **a,** Example transverse slice used for tracer injections. Injections of dextrans were made into the ANC. **b,** Transverse section of an injection of dextran (cyan) demonstrating ipsilateral connections to the cell body layer (CBL). Scale bar: 50 μm. **c,** Transverse section of an injection of dextran (cyan) into the neuropil (NP) demonstrating contralateral connections arising from the CBL. Scale bar: 50 μm. **d,** Horizontal series stained with SMI-31 (cyan), F-actin (magenta) and DAPI (gray) through the ANC from aboral **(i)** to oral **(vi)**. SMI-31 labels a subpopulation of nerve fibers and is useful for illustrating key selected features of fiber architecture. Scale bars: 100 μm**. (i)** Cerebrobrachial tract (CBT) composed of two massive longitudinally running fiber bundles. **(ii)** Brachial territory of the ANC. SMI-31 brachial nerves branch in proximal and distal directions, pooling over segments. **(iii)** Interface between the brachial territory and the sucker territory of the ANC. This territory is dominated by longitudinally running tracts. **(iv-vi)** Sucker territory of the ANC, progressively showing **(iv)** clear contralateral connections and links to adjoining suckers, **(v)** more restricted intermediate ipsilateral connections, and **(vi)** highly local ipsilateral connections. **e,** Maximum projection of a longitudinally cut whole mount labeled with acTUBA (cyan). The oral nerves originate from a sucker enlargement and directly target the corresponding sucker. Scale bar: 500 μm. **f,** Maximum projection of a whole mount of a sucker labeled with acTUBA. The oral nerves can be traced to the sensory epithelium lining the sucker rim. Scale bar: 500 μm. **g,** Oral nerves false-colored in their position in the sensory epithelium. Neighboring nerve fibers target adjoining territories along the sucker rim. Scale bar: 100 μm. **h,** Maximum projection of a horizontal slice through a sucker whole mount labeled with acTUBA (cyan). The sensory epithelium is evenly and densely innervated. Scale bar: 100 μm.

**Supplemental Figure 7:**
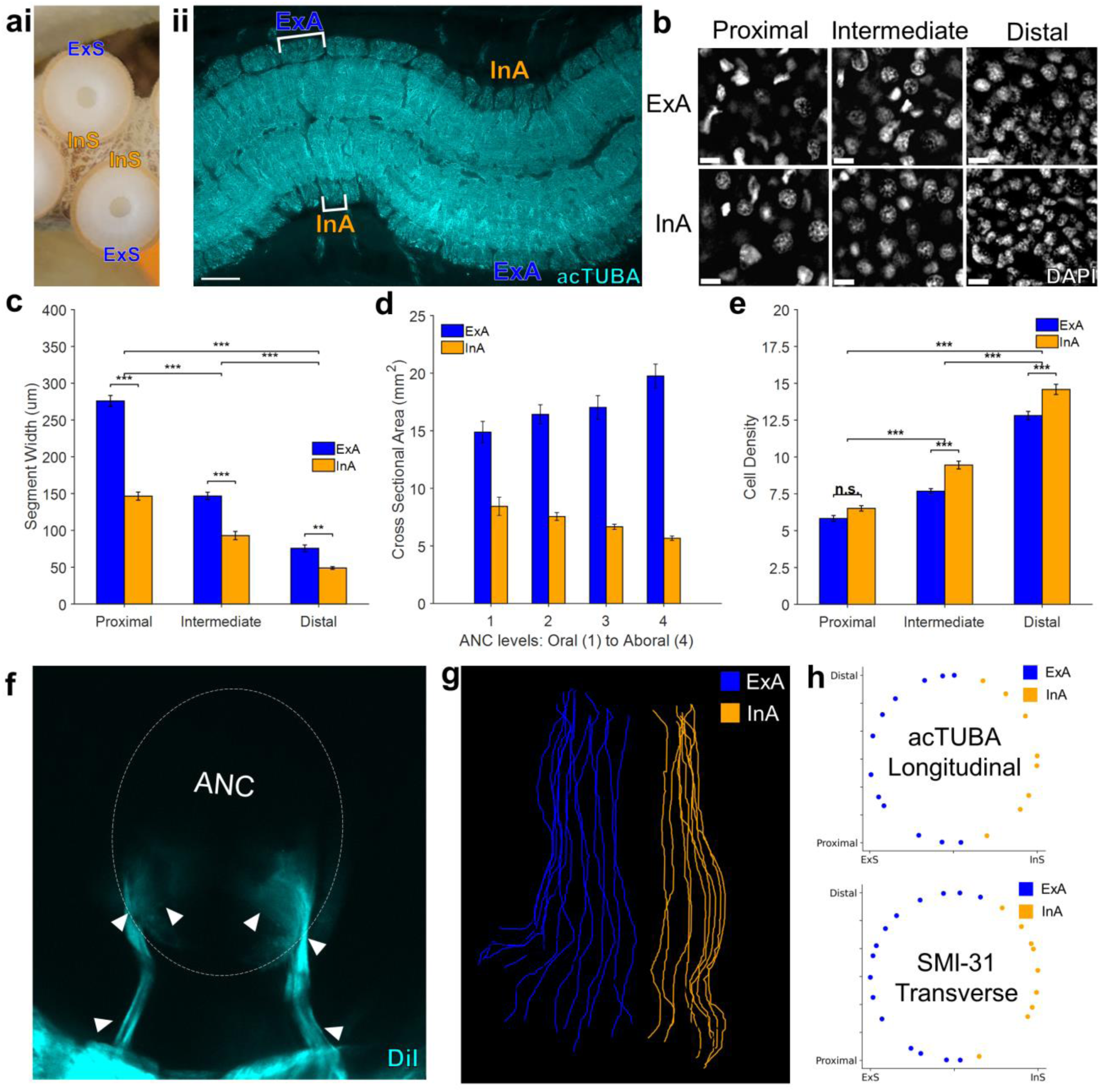
External and internal sides of the ANC. a, (i) The sucker can be divided into the internal side (InS), which hugs the midline of the arm, and the external side (ExS). **(ii)** Horizontal section through the ANC stained with acTUBA (cyan). One side of the ANC (ExA) overlies to sucker’s ExS, and the other (InA) corresponds to InS. Scale bar: 100 μm. **b,** Example patches of DAPI labeling in the ANC used for cell density calculations. Scale bar: 10 μm. **c,** Width measurements of Arm 1 ANC CBLs split into ExA (blue) and InA (orange). Segments on ExA were significantly wider than segments on InA. n = 24 per condition, error bars +/- sem, **p<0.01, ***p<0.001. **d,** Cross-sectional area of segments taken from four horizontal sections stained with acTUBA and spanning the oral to aboral extent of the ANC. Segments corresponding to ExA (blue) had larger cross-sectional areas compared to InA segments (orange). n = 9 per condition, error bars +/- sem. **e,** Cell density calculations. Cell density significantly increases from proximal to distal and is significantly larger for InA segments in the intermediate and distal slices. n = 24 per condition, error bars +/- sem; n.s., not significant, ***p<0.001). **f,** Transverse wholemount with DiI (cyan) crystal placed in a single sucker. The oral nerves enter the ANC on both the InA and ExA. **g,** Traced oral nerve fibers from a transverse whole mount labeled with acTUBA, and tagged for ExA (blue) and InA (orange) targeting. **h,** *Top*- Distribution of oral nerve tips traces from a longitudinal whole mount stained with acTUBA. ExA covered 63%; InA, 37%. *Bottom*-Distribution of oral nerve tips traces from a transverse whole mount stained with SMI-31. ExA covered 64.5%; InA, 35.5%.

**Supplemental Figure 8:**
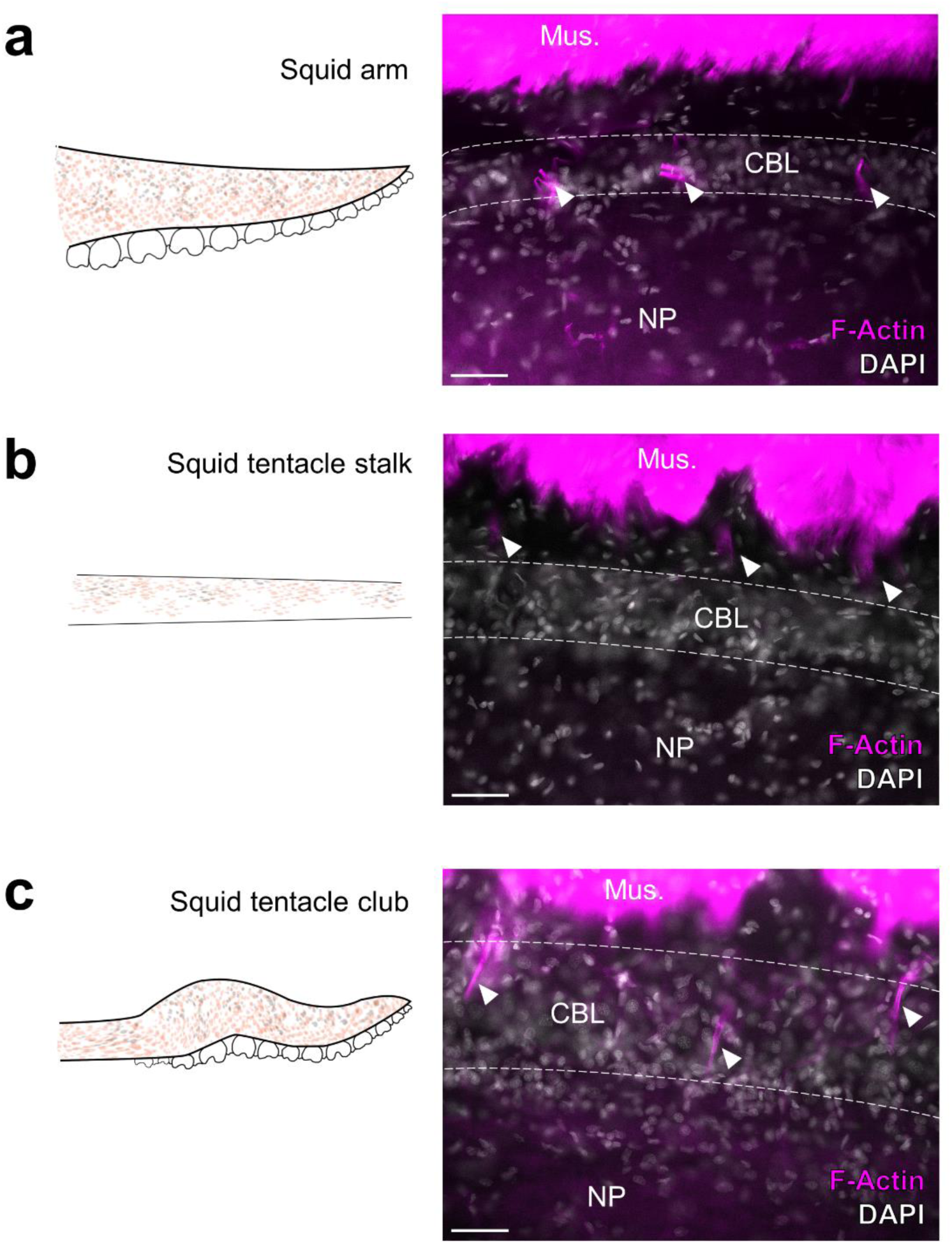
Localization of F-Actin in *D. pealeii* ANC. **a,** Horizonal section through the ANC in *D. pealeii* arm stained for F-actin (magenta) and DAPI (gray). F-actin projections are contained within the CBL. **b,** Horizontal section through the ANC in *D. pealeii* tentacle stalk stained for F-actin (magenta) and DAPI (gray). F-actin projections are beyond the CBL. **c,** Horizontal section through the ANC in *D. pealeii* tentacle club stained for F-actin (magenta) and DAPI (gray). F-actin projections are again within the cell body layer. Scale bars: 100 μm.

